# Rapid, specific detection and quantification of *Yersinia pestis* using a species-specific SNP in the ferric uptake regulator gene (*fur*MAMA)

**DOI:** 10.1101/341321

**Authors:** Pernille Nilsson, Shair Gurbanov, W. Ryan Easterday

## Abstract

*Yersinia pestis*, the causative agent of plague, is responsible for about 700 human cases of bubonic and pneumonic plague each year. Yet the disease is far more prevalent within rodent reservoirs than in humans. One of the main means of outbreak prevention is extensive wildlife surveillance, where accurate and rapid detection is essential to prevent spillover into the human population from which, it may otherwise spread more rapidly and over larger distances. Moreover, detection and quantification of the agent aids in investigative studies to understand aspects of the pathogen such as transmission mechanics, pathology, contamination risk and more. Partially based on a previously developed assay by Gabitzsch et al. 2008 we designed a TaqMan^®^ mismatch amplification mutation assay (TaqMAMA) where a primer leverages a species-specific SNP in the chromosomal single copy ferric uptake regulator gene of *Yersinia pestis*. The assay allows for specific, rapid detection and quantification of *Yersinia pestis* using only a single species-specific marker in a highly conserved virulence gene. This low-cost and simple modification of an existing assay eliminates the need for running multiple molecular markers for pathogen detection or performing time-consuming culturing and counting of colonies for quantification.

## Background

The causative agent of plague, *Yersinia pestis*, reached infamy after causing three devastating human pandemics during recorded history. The disease however, is mainly a vector-borne, wildlife disease, circulating within rodent populations across the world, which occasionally spills over into human populations. Today there are approximately 700 reported human cases of plague worldwide each year with more than 100 deaths (World Health Organization, 2017), most of which occur in Africa. Due to the pandemic history of the pathogen and the continued potential for human outbreaks, close surveillance of its presence in wildlife systems has been incorporated into several local and national surveillance programs, some dating back to the beginning of the 1900s (Melikishvili, 2006).

Plating of infected animal tissues or fleas and subsequent counting of bacterial colonies is a well-established, sensitive and widely used method in many surveillance programs and in diagnostics (Bevins et al., 2012). This is a time-consuming method that exposes the workers to infectious material for extended time. The culturing of *Y. pestis* has historically had challenges due to the relatively slow growth of *Yersinia spp.* compared to the near omnipresent contaminating microbes on traditional media (Dennis et al., 1999; Gupta et al., 2015; Sarovich et al., 2010). To overcome some of these issues, new and improved growth media with increased selectivity for *Y. pestis* have been developed but still struggle with obstacles such as reduced recovery rate of the pathogen and desired level of selectivity leading to underestimation of bacterial numbers (Ber et al., 2003; Riehm et al., 2011; Sarovich et al., 2010). As of today, culturing remains a gold standard for detection and in particular quantification.

The recent development of immunoassay tests for rapid and easy detection in the form of dipsticks have become preferential, particularly for quick diagnostics in challenging field conditions in remote locations (Chanteau et al., 2003; Simon et al., 2013). Yet immunoassays are typically not sensitive enough to pick up cases of low-level infections or early on in infections where only small amounts or no antibody is present (Andrianaivoarimanana et al., 2012; Chanteau et al., 2003; Simon et al., 2013). Other limitations of these assays are potential false negative results due to the lack of detection of strains that have genetic deletions of the assays target (such as the F1 capsular antigen) (Anisimov et al., 2004; Eppinger et al., 2010; Perry and Fetherston, 1997). They are also altogether unsuitable for quantification.

In the era of sequencing and genomics and the power and relative ease of PCR based methods, the gold standard of detection is shifting from culture-based methods towards molecular tools. In this context, PCR based assays are able to detect the pathogen at low quantities making them the most appropriate choice for investigative studies where maximal sensitivity is required. Yet specific detection of bacterial pathogens using PCR or immunoassay tests must often navigate several major obstacles, many being the products of bacterial evolution. Most pathogens evolve out of complexes of related species, where mechanisms like horizontal gene transfer (HGT) allows a constant exchange of genetic elements, often as plasmids or genetic islands (Juhas, 2015). As a result of these evolutionary trends, specific detection of the target of interest, as is the case for many pathogenic bacteria including *Y. pestis and Y. pseudotuberculosis,* can be difficult (Achtman et al., 1999; Easterday et al., 2005; U’Ren et al., 2005). For *Y. pestis,* its recent evolution from *Y. pseudotuberculosis,* has been accomplished through a process of functional gene loss and gain through HGT within *Enterobacteriaceae.* This results in *Y. pestis* being, more or less, a clone of *Y. pseudotuberculosis* with shared elements from other members of *Enterobacteriaceae* (Achtman et al., 1999; Hinnebusch et al., 2016).

Historically, discrimination between *Y. pestis* and *Y. pseudotuberculosis* has been challenging for methods targeting chromosomal sequences, due to the high degree of genetic similarity between the two species (Califf et al., 2015; Chain et al., 2004; Gabitzsch et al., 2008; Neubauer et al., 2000). A work around has been to use multiple assays to detect specific combinations and variants of genes located on the chromosome and/or the plasmids of *Y. pestis* (Stewart et al., 2008; Tomaso et al., 2003). Still, the problems with finding *Y. pestis* specific genes on the main chromosome also extends to plasmid targets, such as the pCD1 plasmid, a homologue of pYV1 found in *Y. pseudotuberculosis* and *Y. enterocolitica* (Hu et al., 1998; Portnoy et al., 1984).

The two most conventional targets for detection of *Y. pestis* both under laboratory and field conditions are the Capsule antigen fraction 1 (*caf1*) gene on the pMT1 plasmid and the plasminogen activator gene (*pla*) on the pPCP plasmid*. Pla* is one of the most widely used markers for PCR based detection of *Y. pestis* in ancient human remains, animals and environmental samples due to it being present on the pPCP plasmid which is present in multiple copies in the bacteria and its presumed high specificity for *Y. pestis* (Harbeck et al., 2013; Stewart et al., 2008; Tomaso et al., 2008). However, recently it has been established that the gene is present and highly conserved in closely related bacterial species, and consequently is no longer considered a specific marker for *Y. pestis* (Armougom et al., 2016; Hänsch et al., 2015). Generally, a minimum of one other independent molecular marker is required to positively determine presence of the bacteria; these include genes *caf1* and Yersinia murine toxin (*ymt*) from the pMT1 plasmid, Low calcium response V antigen (*lcrV*) from the pCD1 plasmid, pesticin gene on pPCP or chromosomal targets such as 16S or Inner membrane protein yihN (*yihN*) gene either in separate or multiplex real-time PCR assays (Iqbal et al., 2000; Stewart et al., 2008; Tomaso et al., 2008; 2003; Woron et al., 2006). Furthermore plasmids are subject to variation in copy number during certain conditions, such as during an infection (Wang et al., 2016). Parkhill et al. showed that *Y. pestis* on average contain about 200 copies of the pPCP1 plasmid (Parkhill et al., 2001), which is partly the reason for the preference of *pla* as target for detection as the gene is present in high copy number, yet the variability in copy number is a severe drawback for quantifying *Y. pestis* concentrations (Stewart et al., 2008; Tomaso et al., 2003).

By combining the established TaqMan^®^ assay with a mismatch amplification mutation assay (TaqMAMA) (Cha et al., 1992), Glaab and Skopek (Glaab and Skopek, 1999) developed a powerful and rapid method of discriminating specific single nucleotide alleles in real-time providing a valuable tool for several fields of science dealing with discrimination between highly similar targets (Achtman et al., 1999; Chain et al., 2004; Lärkeryd et al., 2014). TaqMAMA assays have successfully been designed to specifically amplify, detect and quantify a particular allele when it is paramount to distinguish between highly similar DNA sequences. Most of these have been developed in cancer research (Abbaszadegan et al., 2009; Cha et al., 1992), but also for work in bioforensics or surveillance when positive detection and discrimination of a pathogen from close genetic neighbors is vital (Achtman, 2008; Easterday et al., 2005). Previously, Gabitzsch et al. (Gabitzsch et al., 2008) developed a TaqMan^®^ assay to detect the ferric uptake regulator (*fur*) gene for the specific quantification of *Y. pestis*, yet has become outmoded as new sequences have become available showing the gene is highly conserved in other *Yersinia* and more widely occurs within *Enterobacteriaceae*. Fortuitously, a 100% *Y. pestis*-specific SNP exists within this gene inward from the priming site of the reverse primer (*YpfurR*) from the Gabitzsch assay (Fig. 1). Here we leverage this species-specific SNP to provide a simple and cheap augmentation to the original assay creating the only single target assay for *Y. pestis,* which is quantitative, sensitive and rapid as well as, species-specific.

**FIG 1.**
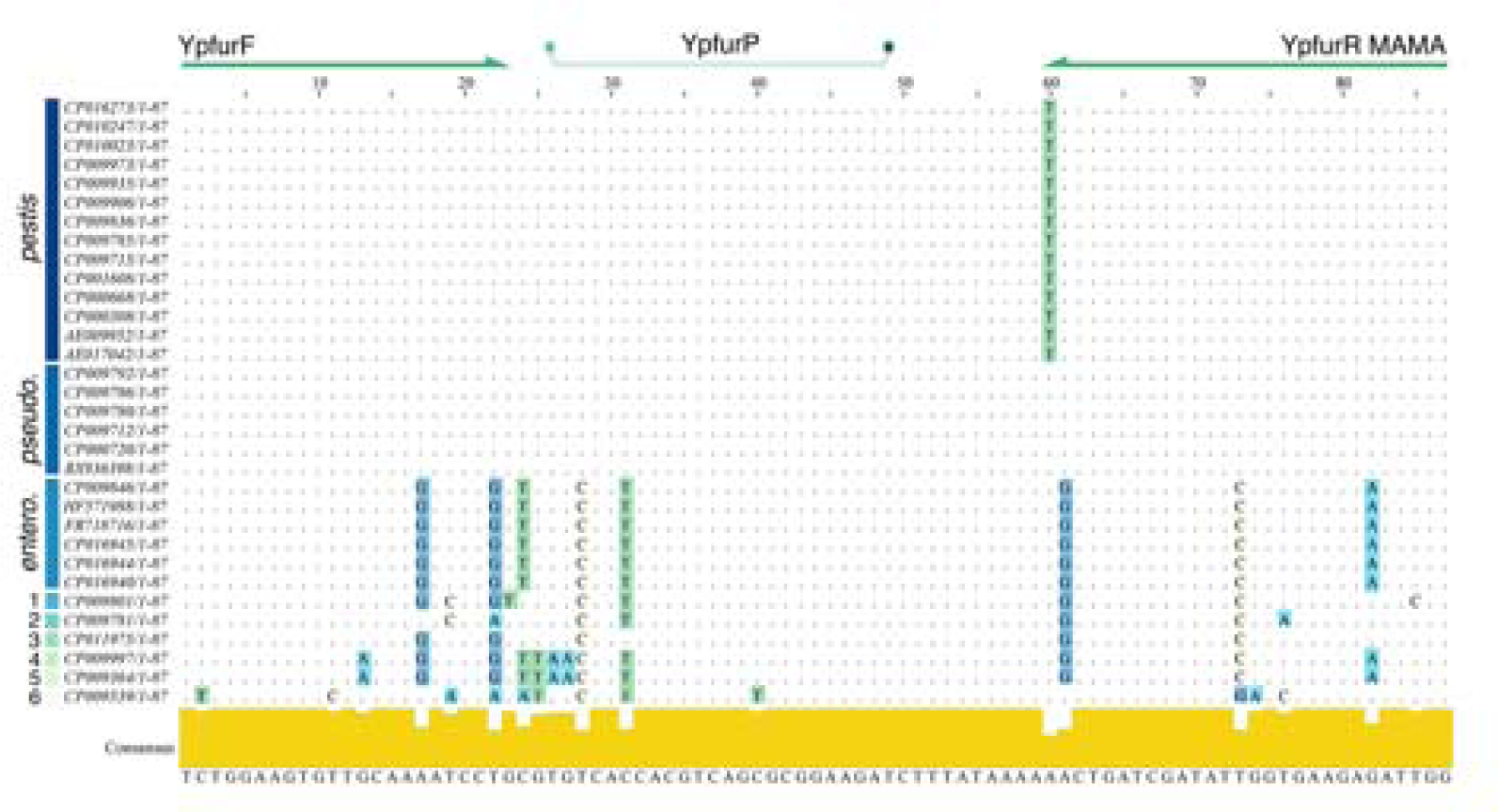
Nucleotide alignment of the 87 bp PCR fragment, targeted by the assay, of the *fur* gene of a representative selection of *Yersinia* spp. Only the nucleotide differences between the strains and the generated consensus is shown. Alignment was generated through an initial BLAST search in Geneious, names (NCBI accession numbers) and sequences were subsequently extracted and entered into Jalview where an alignment was automatically performed to generate the consensus sequence and bar plot. The location of the forward and reverse primers and the probe are indicated on the top of the alignment. *pestis*: a selection of *Y. pestis* sequences covering several biovars and geographic locations, *pseudo.*: a selection of *Y. pesudotuberculosis* sequences, *entero.*: a selection of *Y. enterocolitica* sequences, 1: *Y. intermedia*, 2: *Y. aldovae*, 3: *Y. aleksiciae*, 4: *Y. kristensenii*, 5: *Y. frederiksenii*, 6: *Y. ruckeri*.

## Results

### *In silico* screen of SNP, a private allele for *Y. pestis*

The ubiquitous presence of the *Y. pestis* allele was confirmed by both megaBLAST and BLASTn searches (NCBI) of the *fur* gene sequence from CO92 and the PCR fragment. In both cases the allele was conserved in all 981 published *fur* gene sequences for *Y. pestis* (a global representation), while the alternate allele was present in all 853 available *fur* gene sequences of *Y. pseudotuberculosis* and all 2950 available *fur* gene sequences of the other members of the *Yersinia* genus (May 2018, Fig. 1). A BLASTn search of the *fur* sequence of CO92 against the genome assembly of the strain used in this study, Az-26 (1102) (not published) confirmed the presence of the *Y. pestis* allele in the *fur* gene of this strain, as well as 100% identity with all published *Y. pestis*. The SNP, as to date, is a private allele for *Y. pestis*.

### Specificity

The specificity of the *fur*MAMA assay was assessed by analyzing serial 10-fold dilutions of genomic DNA in quadruplicates from *Y. pestis* and *Y. pseudotuberculosis* alongside the original assay from Gabitzsch et al. (Gabitzsch et al., 2008). We found no difference in the amplification efficiency between *Y. pestis* and *Y. pseudotuberculosis* when using the original reverse primer from the original assay across all DNA concentrations (Fig. 2 (A)). Our augmented assay successfully quantifies concentrations of *Y. pestis* ranging from 5 ng (~100 000 000 genome copies) to 50 fg (~10 genome copies) with a slight shift in amplification efficiency (higher Cq values) compared to when the original reverse primer is used (Fig. 2 (C)). In contrast, we find amplification failure in all dilutions of *Y. pseudotuberculosis* and were only able to achieve false positives using extremely high concentrations of *Y. pseudotuberculosis* template in the *fur*MAMA assay where we saw some weak cross-reactivity in three of four replicates at 5 ng and one of the quadruplicates at 500 pg (Fig. 2 (D)). A combined 90 replicates consisting of 64 non-template controls (NTC), and (assumed) negative tissue and soil samples (from non-endemic areas) did not show any amplification.

**FIG 2.**
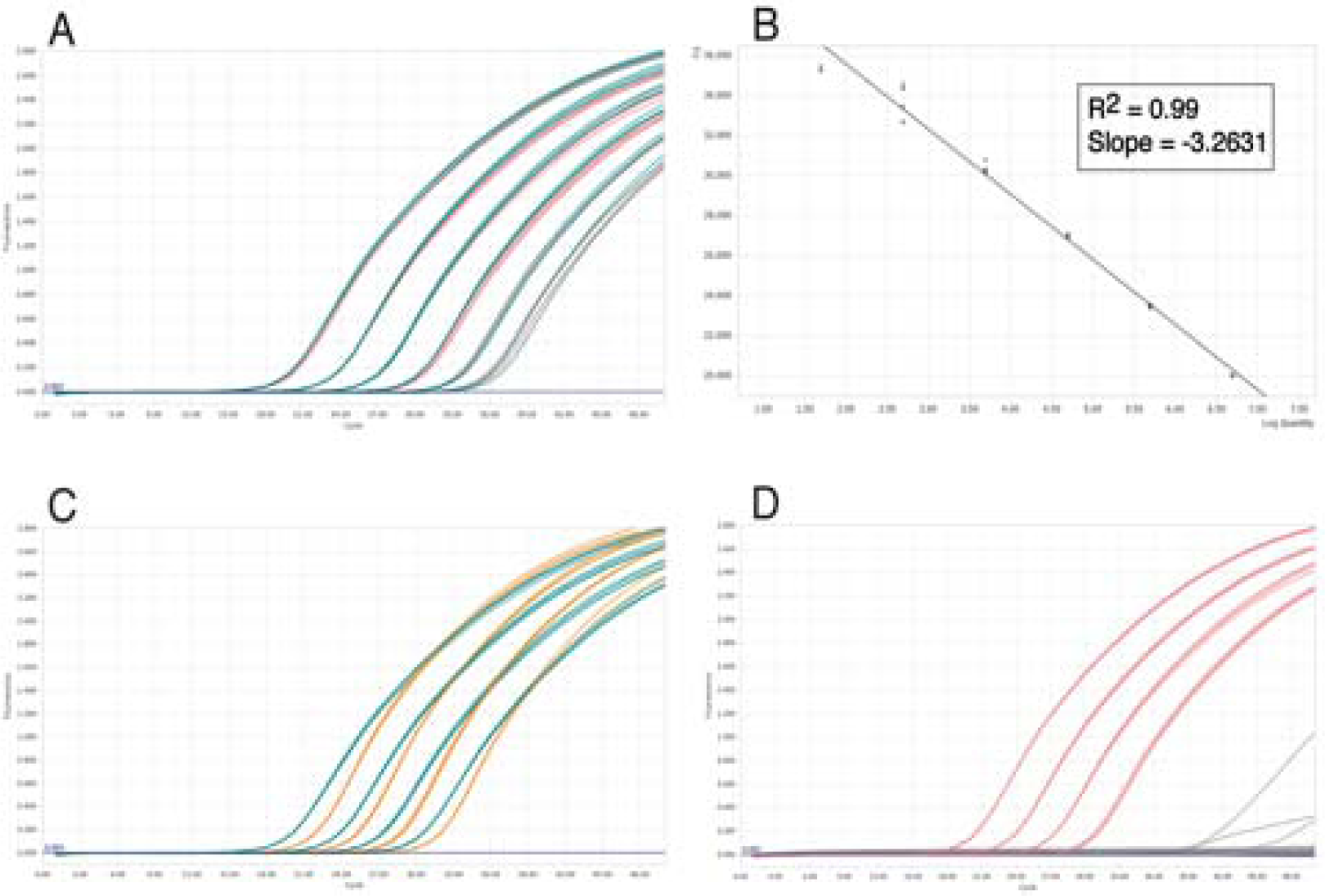
Real-time plots indicating the amplification efficiency and specificity of the original fur assay vs the *fur*MAMA TaqMan^®^ mismatch amplification mutation assay (TaqMAMA). (A) Results of quadruplicate analysis of the 10-fold serial dilutions of *Y. pestis* (blue/teal) and *Y. pseudotuberculosis* (red) DNA (left to right: 5 ng – 50 fg) using the original *fur* assay. (B) The standard curve plot of *Y. pestis* DNA with the *fur*MAMA assay; *R^2^* = 0.99, slope = −3.2631. (C) Results of quadruplicate analysis of the 10-fold serial dilutions of *Y. pestis* with the original fur assay (blue/teal) and the *fur*MAMA assay (orange). Only the first four dilutions are shown. The average Cq values (quadruplicate analysis) in TE buffer were as follows: 5 ng, 17.87; 500 pg, 21.80; 50 pg, 25.33 and 5 pg, 28.64 for the original *fur* assay and 5ng, 19.98; 500 pg, 23.44; 50 pg, 26.96 and 5 pg, 30.36 for the *fur*MAMA assay (curves left to right). (D) Results of quadruplicate analysis of the 10-fold dilutions of *Y. pseudotuberculosis* with the original *fur* assay (red) and the *fur*MAMA assay (grey). Only the first four dilutions are shown. The average Cq values (quadruplicate analysis) in TE buffer were as follows: 5 ng, 18.16; 500 pg, 21.75; 50 pg, 25.23 and 5 pg, 28.96 for the original *fur* assay while the dilutions show overall amplification failure with the *fur*MAMA assay. For the 5 ng *Y. pseudotuberculosis* with *fur*MAMA three of the four replicates amplified with the average Cq value of 40.15. For the 500 pg only one of the quadruplicates amplified and the Cq value was 43.88. No amplification was seen for either the 50 pg or 5 pg dilutions.

### Specific quantification and sensitivity of the assay

To evaluate the ability of specific quantification and the sensitivity of the *fur*MAMA assay, 10-fold serial dilutions of *Y. pestis* were analyzed. The assay successfully amplified *Y. pestis* over a dynamic range of concentrations with a correlation coefficient (R^2^) of 0.99 (Fig. 2 (B)) and was able to consistently quantify as little as 50 fg, which is equivalent to 10 bacterial genomes. The assay showed sporadic amplification at levels of the 5 fg concentration (3 out of 33 reactions).

### Detection by conventional PCR

The *fur*MAMA assay was also run as a standard PCR without probe to test its applicability as a detection only assay for when quantification equipment is not available. The primer pair successfully amplified *Y. pestis* and did not produce a PCR product of the correct size for any of the quadruplicates of *Y. pseudotuberculosis* (results not shown).

### Screening for inhibition by rodent tissue and soil extracts

In order to assess possible effects of foreign DNA on the *fur*MAMA assay we ran target-spiked controls over a standard dilution range of concentrations (5ng to 50fg) into DNAs extracted from rodent tissues and soils. *Y. pestis* DNA was spiked into several types of DNA extracts including ear, tail, liver and spleen from several rodent species including, great gerbil (*Rhombomys opimus*), lab mouse (*Mus musculus*) and bank vole (*Myodes glareolus*) and DNA from soil from a previous study (Turner et al., 2016). Regardless of background DNA (tissue or soil) the assay still detects and quantifies across the range tested without non-specific cross-reactivity. Environmental samples such as soil extracts can show a high degree of inhibition depending on the sample and extraction method used (Whitehouse and Hottel, 2007). Our assay successfully amplified soil extracts spiked with *Y. pestis* although we did observe an approximately 10-fold reduction in amplification efficiency that persisted despite the addition of BSA to the reaction.

## Discussion

Although many PCR-based assays exist for *Y. pestis*, none combines species specificity with accurate quantification. The *fur*MAMA assay is designed to specifically amplify a fragment of the *fur* gene in *Y. pestis* while preventing amplification of the gene fragment in other members of the *Yersinia* genus including its close genetic neighbor *Y. pseudotuberculosis* where the gene fragment only differs by a single SNP. The SNP in the *fur* gene fragment targeted by our *fur*MAMA assay is private to, and ubiquitous in all *Y. pestis*. Specificity is acquired in a single assay through allele-specific amplification of this SNP in a key virulence gene, *fur*, whose presence is found throughout *Enterobacteriaceae* without the need for further molecular targets (Fig. 1). Leveraging a specific SNP within a gene, as done here, is an efficient and robust method for specific detection of a given bacterial pathogen and can aid in pathogen detection in metagenomes using next generation sequencing where close relatives are present in high numbers (Valseth et al., 2017). This circumvents the present issues of specificity when targeting pathogens with high degree of genetic similarity. A recent large-scale comparative study of *Y. pestis* and *Y. pseudotuberculosis* genomes showed that fewer genes than previously reported in smaller studies were unique to *Y. pestis* (Califf et al., 2015). However, both specific combinations, variants of genes and SNPs, the last example demonstrated here, will be unique to this slowly mutating pathogen (Cui et al., 2013).

The original assay published by Gabitzsch et al. (Gabitzsch et al., 2008) was developed to quantify *Y. pestis* in fleas for their work on flea transmission in a laboratory experimental setup and not initially aimed at diagnostics or quantification in field-collected samples. They tested the specificity by running PCR on reference DNA from 28 species, including *E. coli* and other gram-negative bacteria, and reported cross-reactivity for 6 *Yersinia* species, including *Y. enterocolitica*. Surprisingly, they did not report cross-reactivity with *Y. pseudotuberculosis*. BLAST of the derived *fur* gene fragment from the Gabitzsch assay resulted in 99% overall homology with *Y. pseudotuberculosis* sequence and high homology between other *Yersinia* spp. Indeed, we found the original assay amplifies both *Y. pestis* and *Y. pseudotuberculosis* with equal efficiency (Fig. 2 (A)). In contrast, the amplification failure of *Y. pseudotuberculosis* with the YpfurR_MAMA primer establishes our assay’s ability to distinguish between the two highly genetically similar pathogens, as well as the ability to quantify *Y. pestis* over a broad range of concentrations (Fig. 2(D and C)). The *Y. pestis* SNP in the *fur* gene is universally present in all available *Y. pestis* genomes, which limits the probability of false negative results. Conversely, the absence of the mutation in all other *Yersinia* and *Enterobacteriaceae* reduce the likelihood of false positive results. Our assay presented false positives and very low-level cross-reactivity at the highest concentration of *Y. pseudotuberculosis* DNA produced from whole genome amplification (WGA). The lack of access to a larger collection of *Y. pseudotuberculosis* prohibited us from further testing of cross-reactivity but based on the specificity testing *in silico* and by targeting a conserved SNP in the *Y. pestis* genome we are confident this will not be a major problem.

## Conclusions

Our assay provides an alternative to the longstanding culturing method and the immunoassays by providing a highly sensitive and rapid way of both specifically detecting and quantifying *Y. pestis* in tissue samples. To our knowledge, this is the first assay to target a species-specific SNP in a chromosomal gene of *Y. pestis* with the aim of detecting and quantifying the amount of the bacteria in tissue samples.

## Materials and Methods

### Bacterial strains and DNA extractions

*Y. pestis* strain Az-26 (1102) was isolated from an organ suspension of *Meriones vinogradovi* in 1969, 1.5 km southeast of Sirab village in the Nakhchivan Autonomous Republic of Azerbaijan. The culture has been stored at the Republican Anti-Plague Station laboratory in Baku, Azerbaijan. The culture became non-viable, yet it was still possible to isolate DNA from the material in 2012 using Qiagen DNeasy^®^ Blood and Tissue kit following extraction protocol for gram-negative bacteria.

*Y. pseudotuberculosis* Type III strain was donated by Jack C. Leo and Dirk Linke (University of Oslo). DNA was isolated from broth culture using Qiagen Blood and Tissue kit following manufacturer’s instructions (Qiagen Inc., USA).

### Rodent DNA

No animals were killed for the purpose of this study. We procured samples of rodent DNA for this study by reaching out to both internal and external colleagues with existing DNA samples stored. The DNA samples stem from unpublished work as detailed below.

The great gerbil DNA was provided by colleagues in China where the male individual was captured as part of work carried out in the Junggar Basin of Xinjiang Province, China (unpublished).

The C57BL6 mouse DNA stem from work conducted at the Institute of Immunology at Oslo University Hospital, Rikshospitalet, Norway (unpublished). The animal was originally obtained from Janvier labs (https://www.janvier-labs.com/rodent-research-models-services/researchmodels/per-species/inbred-mice/product/c57bl6jrj.html).

The bank vole DNA belongs to the EcoTick-project on tick-/rodent-borne diseases conducted at the University of Oslo (unpublished).

### Whole genome amplification (WGA) and standard curves

Whole genome amplification was performed on both bacterial strains using the REPLI-g Mini Kit (Qiagen) following the manufacturer’s instructions for subsequent creation of a dynamic range of DNA starting at high concentrations for standard curves. DNA concentrations after WGA were determined using the Qubit dsDNA BR Assay kit (molecular probes, Life Technologies) and the Qubit 2.0 fluorometer. The standard curves were generated from that starting concentration as serial 10-fold dilutions in TE buffer in the following concentrations: 5 ng, 500 pg, 50 pg, 5 pg, 500 fg, 50 fg and 5 fg.

### TaqMAMA Primers and Probe design

Forward primer and probe were identical to oligos designed by Gabitzsch et al. (Gabitzsch et al., 2008) while the reverse primer was modified to a MAMA primer to specifically amplify *Y. pestis.* This required the ultimate 3’ base to be complementary to the *Y. pestis* SNP while the penultimate 3’ base was designed to mismatch with the shared sequence between *Y. pestis* and *Y. pseudotuberculosis* (underlined in Table 2). Several reverse MAMA primers were designed with different nucleotide mismatches at the 3’ penultimate position. These were tested for specificity and amplification efficiency before ultimately choosing the YpfurR_MAMA primer.

**Table 1.**
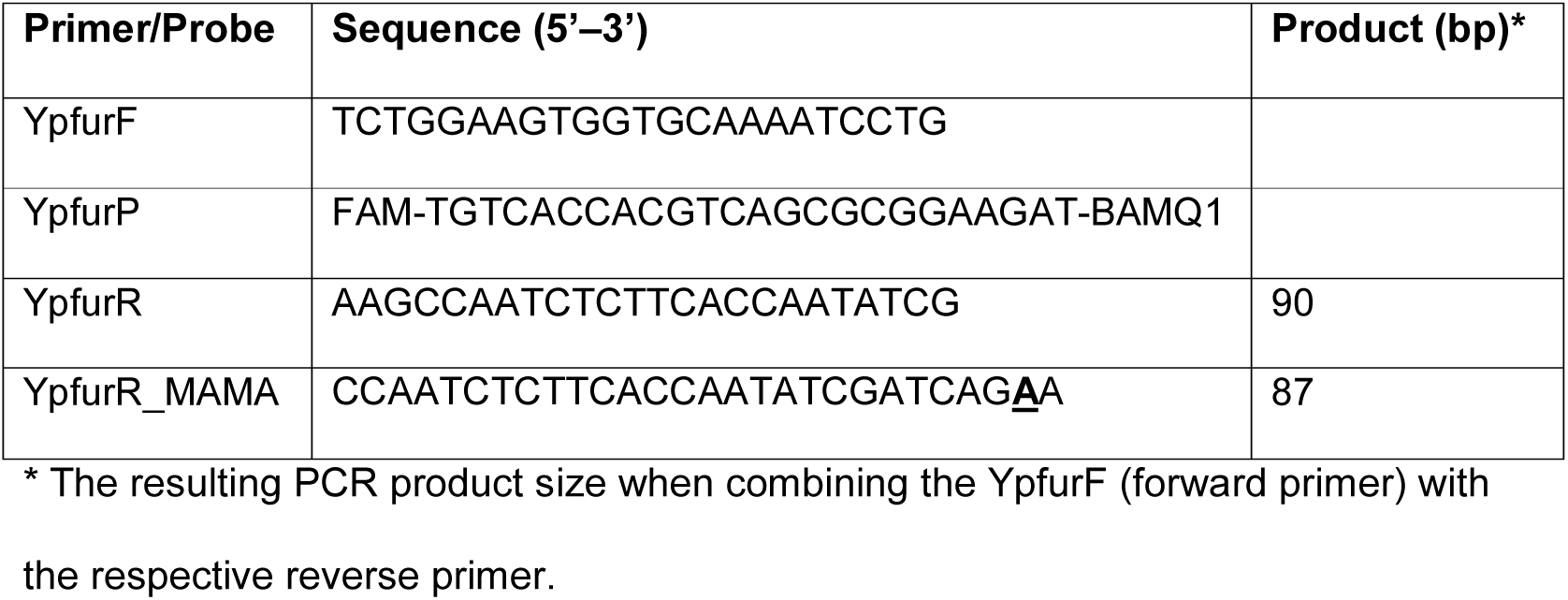
List of Primers and probe sequences assayed in qPCR and PCR reactions

### TaqMAMA PCR protocol

For all experiments, PCR was conducted in 96-well plates (Roche LightCycler^®^ 480 Multiwell Plate 96) using 10-μL reactions that contained final concentrations of 600 nM of both forward and reverse primers, 250 nM probe, 1x Express qPCR Supermix (Invitrogen, by Life Technologies)) and 1μL template DNA. Real-time PCR was performed on a LightCycler^®^ 96 Instrument (Roche) as a two-step PCR with the following conditions: 95 °C for 2 min and 50 cycles of 95 °C for 15 s and 64 °C for 60 s.

### Specificity tests

Given limited access to a large strain collection of *Yersinia*, specificity was largely assessed using bioinformatics (BLAST) of available sequence data (see “Bioinformatics” below). Therefore, the specificity and kinetics of the assay was done through analyzing genomic DNA from a representative of both alleles (see Fig. 1); *Y. pestis* strain Az-26 (1102) and *Y. pseudotuberculosis* Type III strain running both original reverse primer (YpfurR) and the modified MAMA primer (YpfurR_MAMA) in separate, parallel reactions over a dynamic range of DNA concentrations (5 ng to 5 fg).

### Sensitivity tests

Sensitivity of the furMAMA assay was determined by analyzing serial 10-fold dilutions of genomic DNA from the *Y. pestis* strain Az-26 (1102) ranging from 5 ng to 5 fg with the lowest dilution being equivalent to one bacterial genome.

### Testing applicability on tissue and environmental samples

The applicability of the assay on rodent tissue samples was tested by running, in parallel, DNA extractions of liver and spleen from a great gerbil (*Rhombomys opimus*), DNA from the tail of a C57BL6 mouse (*Mus musculus*) and DNA extracted from the ear of a Bank vole *(Myodes glareolus*) with and without spike-in of *Y. pestis* DNA standard curve. Real-time PCR was conducted in 96-well plates (Roche LightCycler^®^ 480 Multi-well Plate 96) using 10-μL reactions that contained final concentrations of 600 nM of both forward and reverse primer, 250 nM probe, 1x Express qPCR Supermix (Invitrogen, by Life Technologies)), 1 μL template DNA and for the spiked samples an additional 1 μL DNA from one of the standard curve concentrations.

The ability of the assay to detect *Y. pestis* despite the inhibitory effects often seen in environmental samples, was tested by using DNA extractions of soil samples diluted 1:10. The soil samples were run in parallel with non-acetylated Bovine Serum Albumin (New England Biolabs) added, with or without DNA from the *Y. pestis* standard curve in the following manner 1) soil + BSA, 2) soil + standard curve and 3) soil + standard curve + BSA. Real-time PCR was conducted as above with the exception that the reactions containing BSA did so in a final concentration of 1mg/ml.

### Detection using standard PCR

The primers were also tested with standard PCR to determine their applicability as a pure detection-based assay. PCR was conducted in 8-well PCR strips in 10-μL reactions that contained a final concentration of 600 nM both forward and reverse primer, 1x Express qPCR Supermix and 1 μL template DNA. PCR was performed on a MJ Research PTC-200 Peltier Thermal Cycler Instrument as a two-step PCR with the following conditions: 94 °C for 2 min and 40 cycles of 94 °C for 15 s and 64 °C for 60 s. Successful amplification was confirmed by running a 3% agarose gel stained with GelRed (Biotium) at 75 V for 1 hr and 20 min in TAE buffer.

### Bioinformatics

Specificity of the SNP was assessed across available *Yersinia* and other related *Enterobacteriaceae* genomes by performing both megaBLAST and BLASTn searches of the PCR fragment and the whole *fur* gene from *Y. pestis* CO92 (Accession no. NC_003143.1, locus_tag=YPO2634) as query in the NCBI database using default parameters. BLAST hits were inspected to confirm which allele was present in the sequence. Finally, the presence of the *Y. pestis* allele in the strain used in this study (Az-26 (1102)) was confirmed by a BLASTn search using the *fur* gene from *Y. pestis* CO92 as query against a BLAST database generated from the Az-26 (1102) genome assembly. From these data an alignment was made using representative sequences from BLAST searches through Geneious (https://www.geneious.com) and aligned using Jalview (Waterhouse et al., 2009).

## Abbreviations

Fur gene: Ferric uptake regulator
BSA: Bovine serum albumin
Caf1: Capsule antigen fraction 1
Cq: Quantification cycle
HGT: Horizontal gene transfer
lcrV: Low calcium response V antigen
Pla: Plasminogen activator
SNP: Single nucleotide polymorphism
TAE: Tris-acetate-EDTA
TaqMAMA: TaqMan^®^ mismatch amplification mutation assay
TE: Tris-EDTA
WGA: Whole genome amplification
yihN: Inner membrane protein yihN
Ymt: Yersinia murine toxin

## Declarations

### Acknowledgements

We would like to thank Jack C. Leo and Dirk Linke at the Department of Biosciences, University of Oslo for providing the *Y. pseudotuberculosis* Type III strain. For the use of the great gerbil DNA, the C57BL6 mouse DNA and the bank vole DNA we would like to thank Ruifu Yang (Beijing Institute of Microbiology and Epidemiology, China) and Yujiang Zhang (Xinjiang CDC, China), Shuo-Wang Qiao (Institute of Immunology at Oslo University Hospital, Rikshospitalet, Norway) and Atle Mysterud (University of Oslo, Norway). We would also like to thank Bradd Haley for bioinformatics support. Lastly, we would like to thank Monica H. Solbakken and Boris V. Schmid for helpful comments on the manuscript.

### Authors contributions

WRE and PN conceived and planned the experiments and analyzed the data. PN carried out the experiments and performed bioinformatics. SG procured and isolated the *Y. pestis* strain Az-26 (1102) for this study. The manuscript was written with input from all authors. All authors read and approved the final manuscript.

### Availability of data and materials

The *Y. pestis* strain analyzed during this study (Az-26(1102)) is not publicly available due to said strain being non-viable at isolation. The genome sequence is available from the corresponding author upon reasonable request.

### Consent for publication

Not applicable.

### Ethics approval and consent to participate

The DNA samples used in this study stems from existing samples stored by internal and external colleagues and hence the appropriate approvals were previously obtained as specified below.

The use of great gerbil tissue was approved by the Committee for Animal Welfares of Xinjiang CDC, China.

The bank vole was captured as part of EcoTick-project on tick-/rodent-borne diseases led by Atle Mysterud (University of Oslo) and permission for trapping rodents were given by the Norwegian Environmental Agency (ref 2013/11201) and hence conform to the Norwegian laws and regulations.

The C57BL6 mouse was used in a project at the Institute of Immunology at Oslo University Hospital, Rikshospitalet, Norway and is subject to protocol number 3775 from the Norwegian Food Safety Authority. This agency evaluates all applications and approvals for ethical handling of animals and animal experiments in Norway.

